# Inducible degron-dependent depletion of the RNA polymerase I associated factor PAF53 demonstrates it is essential for cell growth and allows for the analysis of functional domains

**DOI:** 10.1101/780361

**Authors:** Rachel McNamar, Zakaria Abu-Adas, Katrina Rothblum, Lawrence I. Rothblum

**Author notes:** To whom correspondence should be addressed: Lawrence Rothblum, Department of Cell Biology, BMSB 553, University of Oklahoma College of Medicine, Oklahoma City, OK. 73104.

## Abstract

Our knowledge of the mechanism of rDNA transcription has benefitted from the combined application of genetic techniques in yeast, and progress on the biochemistry of the various components of yeast rDNA transcription. Nomura’s laboratory derived a system in yeast for screening for mutants essential for ribosome biogenesis. Such systems have allowed investigators to not only determine if a gene was essential, but to analyze domains of the proteins for different functions in rDNA transcription *in vivo*. However, because there are significant differences in both the structures and components of the transcription apparatus and the patterns of regulation between mammals and yeast, there are significant deficits in our understanding of mammalian rDNA transcription. We have developed a system combining CRISPR/Cas9 and an inducible degron that allows us to combine a “genetics-like” approach to studying mammalian rDNA transcription with biochemistry. Using this system, we show that the mammalian homologue of yeast A49, PAF53, is required for rDNA transcription and mitotic growth. Further, we have been able to study the domains of the protein required for activity. We have found that while the C-terminal, DNA-binding domain (tWH) was necessary for complete function, the heterodimerization and linker domains were also essential. Analysis of the linker identified a putative DNA-binding domain. We have confirmed that the helix-turn-helix (HTH) of the linker constitutes a second DNA-binding domain within PAF53 and that the HTH is essential for PAF53 function.

---

Ribosome biogenesis is essential for homeostasis and cell growth, mitotic or hypertrophic. This complex process is energetically expensive, consuming up to 60% of a yeast cell’s RNA synthetic capacity (1,2). As such, it is subject to multiple levels of regulation. The rate-limiting step in this complex process is transcription of the pre-ribosomal RNA (rRNA) genes (rDNA) by RNA polymerase I (Pol I)(3–5). Abnormalities in the rates of rRNA synthesis and ribosome biogenesis cause a broad range of human diseases, the ribosomopathies, which can affect individuals in ontogenetic and tissue specific patterns (6–10). Furthermore, the pleomorphic nucleoli of tumor cells, the sites of rDNA transcription, are characteristic of neoplasia and reflect dysregulation of rRNA synthesis in cancer (11–15). rDNA transcription requires a unique set of transcription factors not used by the other nuclear polymerases, and we are just beginning to understand the biochemistry and functional interactions of these mammalian factors. Many of the genes whose protein products are involved in ribosome biogenesis were identified in CRISPR screenings of essential genes (16–19). High-throughput screening of mammalian cells for proteins that affect nucleolar number has recently revealed proteins involved in regulating ribosome biogenesis (20,21). Such procedures pave the way for the exploration of pathways/proteins involved in ribosome biogenesis and human disease. However, it does not provide for biochemical analysis of function.

Nomura’s laboratory demonstrated that the synthesis of the pre-rRNA transcript is the only essential function of Pol I (22). This provided a screen for mutants dependent on Pol II-driven synthesis of rRNA from the GAL promoter (22,23). These “*rrn*” mutants were defective in ribosome biogenesis *in vivo*; confirmed by the observation that mutations in genes encoding unique subunits of Pol I were isolated (23,24), as well as genes encoding other components of the committed template. A second important advance in the study of yeast Pol I was the development of an *in vitro* transcription system using a crude extract and the fractionation of those extracts and purification of defined transcription complexes (25–29). The combination of the genetic and biochemical approaches significantly enhanced our understanding of the mechanism and regulation of rDNA transcription in yeast.

A fully functional molecule of yeast RNA polymerase I consists of 15 subunits. This total includes the core Pol I, a heterodimer of RPA49-RPA34.5 and RRN3. Five of the core subunits are shared with the other two polymerases and two are shared with Pol III (30,31).

Two of the Pol I subunits, yeast A34.5p (A34) and A49p (A49) form a heterodimer with still poorly defined roles in rDNA transcription. The heterodimer of RPA49/RPA34.5 is easily dissociable from the polymerase, and the association of the mammalian homologs PAF53/PAF49 with Pol I is subject to growth-related regulation (32–35).

While *S. cerevisiae* A49 is not *essential* for viability (36), deletion of yeast A49 results in colonies that grow at 6% of the wild type rate at 25C (36). Similarly, when the *S. pombe* homologue, RPA51, was deleted (37), specific rDNA transcription was reduced 70% (no effect on nonspecific polymerase activity), cells failed to grow at 25C and grew at half the wild-type rate at 30C. Deletion of the other partner in the heterodimer, A34, has a minor effect on growth or rRNA synthesis, but results in a polymerase that loses the A49 subunit upon purification (38). Biochemical purification of Pol I results in two fractions, one of which does not contain either A49 or A34.5 (39). This is complemented by the observation that a large fraction of the polymerase particles prepared for cryo-EM are free Pol I enzymes that “either lacked the A49/A34.5 subcomplex or displayed a flexible clamp-stalk region”(40). Interestingly, most of the interactions between the heterodimer and Pol I in yeast appear to be mediated by the A49 subunit (41). The N-termini of A34 and A49 are required for heterodimerization. The heterodimer stimulates polymerase nuclease activity and has a triple *β*-barrel domain similar to the core of TFIIF and the Pol III heterodimer of C37/C53 (42–44). The C-terminus of yeast A49 contains a domain with dual winged helices (tandem winged helix, t-WH) (42) that is capable of DNA-binding and resembles a similar element in TFIIE (42) and C34 (45,46). Mutations within the tWH of A49 result in increased sensitivity to 6-azauracil and mycophenolic acid and defects in transcription elongation (47). However, these same mutants demonstrate reduced Pol I occupancy at the rDNA locus and lower levels of recruitment of Pol I and Rrn3 at the promoter (47). This would suggest that the mutations may have affected the binding and release of Rrn3 from Pol I as would be necessary for initiation complex formation and transition to the elongating state (36).

Mammalian cells contain homologues to yeast A49 and A34. Muramatsu’s laboratory identified two polymerase-associated factors referred to as PAF53, the homologue to A49, and PAF49, the homologue of A34. They also reported they were essential for promoter-specific transcription (32,33). Biochemical and biological experiments indicate that PAF49 and PAF53 are associated with the active form of Pol I. Yamamoto *et al*. reported that PAF53 and PAF49 were associated with a fraction of the core Pol I molecules (32). Hannan *et al*. confirmed this observation and estimated that 60% of the polymerase molecules in a rat hepatoma cell line contained PAF53 (48). Yamamoto *et al*. also reported on the growth-dependent nucleolar localization of PAF49, a result subsequently confirmed by our laboratory (34,35,49). Most recently, several CRISPR/Cas9 based screenings of the mammalian genome identified PAF53 and PAF49 as being “essential” genes (16,17), which was confirmed when we found that we could not isolate cell lines that failed to express PAF53 (50).

Results from our laboratory and others establish that the assembly of Pol I-specific “**p**olymerase **a**ssociated **f**actors” (Rrn3 and the heterodimer of PAF49 and PAF53) with Pol I is a necessary step in rDNA transcription (25,28,32,33,37,47,49,51–74). While genetic studies in yeast and KO studies in mammalian cells demonstrate that the PAF53/PAF49 complex is essential for cellular physiology (17,42,50,75), their roles in rDNA transcription are unknown. Moreover, because there are significant differences in both the structures and components of the transcription apparatus and the patterns of regulation between mammals and yeast, studies on yeast rDNA transcription leave significant holes in our understanding of mammalian rDNA transcription. As transcription by Pol I is essential for cell growth or viability (76–79), it has been considered impossible to generate cell lines in which the essential genes are knocked out and maintain the population (17,50). While shRNA and siRNA will knockdown the levels of the proteins, this can take days, depending upon the half-life of the protein, and allows compensatory mechanisms to occur. To facilitate the study of rDNA transcription in mammalian cells, we sought to develop a system that would allow us to rapidly knockdown the protein levels and replace them with mutants. This system would enable us to combine a “genetics-like” approach to studying mammalian rDNA transcription with biochemistry. Using this system, we have found that PAF53 is required for rDNA transcription and mitotic growth and we have begun to identify domains of the protein involved in rDNA transcription.

Auxins, such as indole-3-acetic acid (IAA), promote the interaction between SCF^TIR1^ (TIR1), an auxin-dependent E3 ubiquitin ligase [containing, SKP1, Cullin, and an auxin-dependent F-box (80–84)] and the auxin or Indole-3-Acetic Acid (AUX/IAA) family of transcription factors (85,86). TIR1 is the plant-specific, auxin-dependent, F-box protein ***t***ransport ***i***nhibitor ***r***esponse 1 protein. In the presence of auxin, TIR1 associates with the IAA transcription factor. This induces the rapid ubiquitination and subsequent proteasome-dependent degradation of the AUX/IAA family of transcription repressors that contain an auxin inducible degron (AID). Several laboratories have demonstrated that the degron of the IAA transcription repressor can be used to target proteins for degradation. The AID from the IAA17 protein of *Arabidopsis thaliana* has been used to program various proteins for rapid degradation in yeast and animal cells (80,81,87).

We investigated the use of that system to study transcription by Pol I in mammalian cells. In particular, we have focused on PAF53. In our studies, we have examined the use of different forms (lengths) of an auxin inducible degron. This included the degron found in auxin-responsive protein IAA17 (IAA17, indole-3-acetic acid inducible 17) of *Arabidopsis thaliana*, aa 1-229 and parts thereof including aa 68-111, as well as aa 82-94, the 13 amino acid conserved core of the degron, QVVGWPPVRSYRK (82). Further, we found there was no requirement for positioning the degron either N- or C-terminal confirming the finding reported by Nishimura *et al*. that there was no specific requirement for positioning. (80). Also, the degron worked when it was internal on the target protein. Finally, we determined that the optimal degradation of nucleolar proteins required that we add a nuclear localization signal (NLS) to the TIR1 from *Oryza sativa* (81).

Using CRISPR/Cas9, we tagged the C-terminus of the endogenous PAF53 genes with a 43 amino acid, auxin inducible degron (AID) from IAA17 (aa 68-111) using homology directed repair and a donor fragment with short homology arms (~125 bp). Recombination in the PAF53 gene was confirmed by western blot analysis of the expressed protein and PCR analysis of the “new” gene. We found that the degron functioned to target the protein for degradation in HEK293 cells that constitutively expressed NLS-tagged TIR1. When cell lines that expressed PAF53-AID were treated with IAA, the PAF53-AID was eliminated within one hour. This results in the inhibition of rDNA transcription and causes cell cycle arrest. Using cell cycle arrest as the phenotype, we subsequently investigated the domains of PAF53 required for cell division. *In silico* analysis of mammalian PAF53 revealed structures strikingly similar to that of yeast A49. These included the N-terminal dimerization domain, a linker and the C-terminal tandem winged helix. We have previously confirmed the functional homology of the N-terminal dimerization domain (88), and we now demonstrate that the C-terminal tWH has DNA-binding activity similar to that of the yeast domain. We found that deletion of the N-terminal dimerization domain significantly inhibited the ability of ectopic PAF53 to rescue cell division. Interestingly, deletion of the C-terminal tWH had an intermediate effect on rescue, while further deletion of the linker inactivated PAF53. This suggests that both DNA-binding and the ability to dimerize with PAF49 are required for rescue. Most significantly, our data demonstrate that the linker itself must have a function. We identified a DNA-binding motif in the linker and found that DNA-binding by the linker was required for PAF53 activity. Finally, our data imply that the binding of PAF53 to Pol I requires dimerization with PAF49 a result that would be consistent with our previous observation that PAF49 could bind to Pol I independently of PAF53, and very different from that found in yeast.

## Results

Our first experiments were designed to determine whether a nucleolar protein could be targeted for degradation by TIR1. As shown in Figure 1, we found that when amino acids 1-229 of *A. thaliana* IAA17 were linked to H2B tagged with YFP and an AID (Panel B, Figure 1), the protein was degraded when HEK293 cells were cotransfected with a vector expressing TIR1 and treated with IAA. Subsequently, we found that the same AID could target PAF53 expressed transiently (Panel C, Figure 1). We also found that the 43 amino acid degron (aa 68-111) of IAA17 would drive auxin-dependent degradation in HEK293 cells (Panel C, Figure 1). We then examined the possibility of using an Aux/IAA 13–amino acid domain II consensus sequence, aa 82-94, that has been shown to work in plants (82). That domain did not work in our cells (Lanes 5 and 6, Panel C, Figure 1). These experiments also serve to demonstrate that activation of TIR1 by IAA did not target the endogenous PAF53 (lower band in Panels C and D). Subsequently, we have used the 43 amino acid degron as the AID in our constructs. Nishimura *et al*. (80) reported that the AID tag could be on either the N- or C-termini of proteins and still support auxin induced degradation. As shown in Panel D, Figure 1, we confirmed that observation. In mammalian cells expressing TIR1, the AID can be on either terminus of the target protein, and as shown below, it can also be internal.

**Figure 1.**
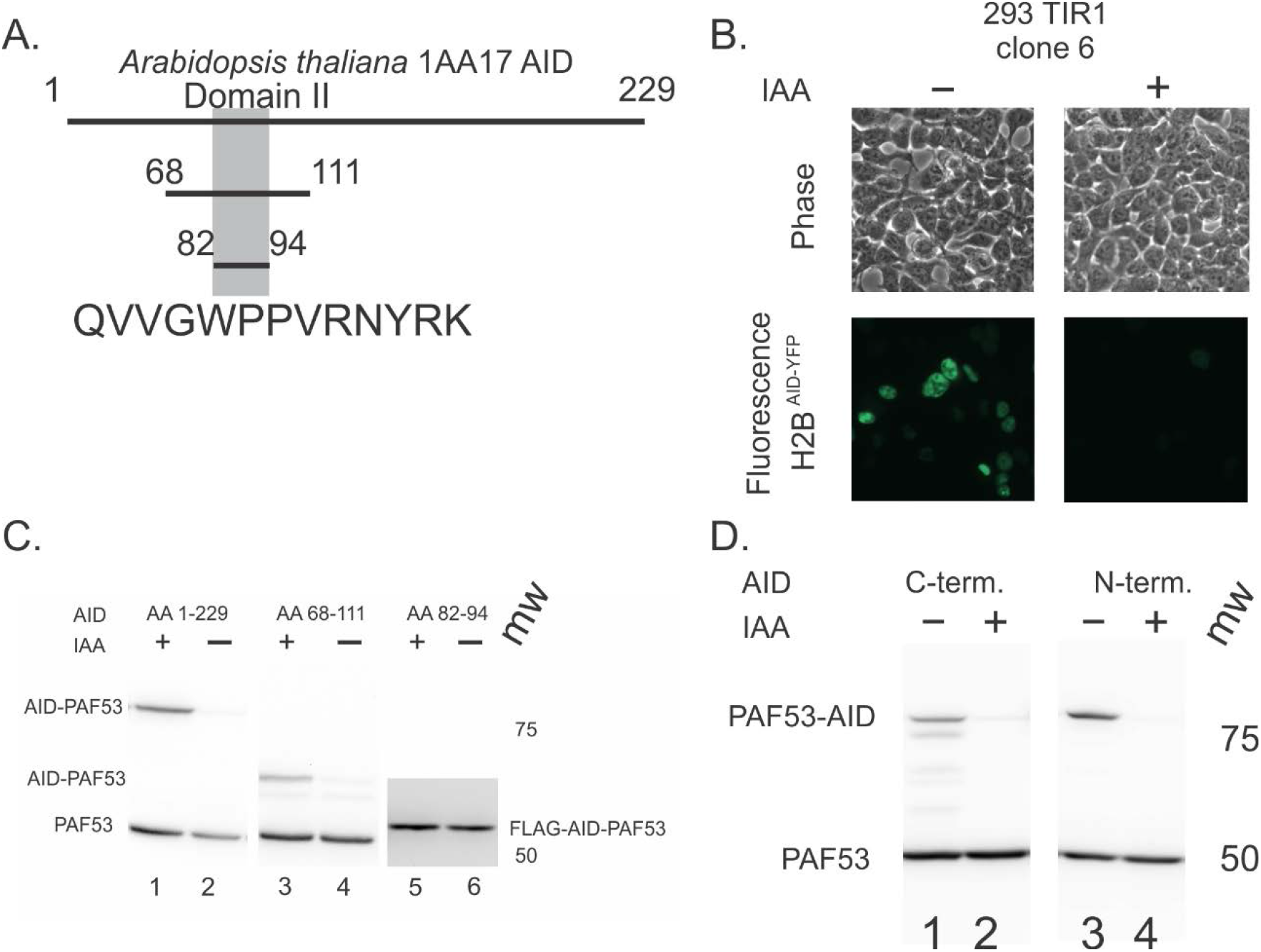
The AID of Auxin/IAA17 of *A. thaliani* imparts auxin-dependent degradation to nucleolar proteins. A. Schematic of IAA17 and the regions used to transfer auxin-dependent degradation. The consensus sequence from Domain II is shown. B. H2B-GFP-IAA17 is targeted for auxin-dependent degradation in HEK293 cells that express TIR1. HEK293 cells constitutively expressing TIR1 were transfected with a plasmid (pcDNA3.1) coding for H2B-GFP-IAA17 for 24 hr, and then treated with 500 *μ*M indole acetic acid (IAA) in water. Amino acids 1-229 of *A. thaliana* IAA17 were linked to H2B tagged with YFP and an AID. C. Amino acids 1-229 and 68-111 can program PAF53 for auxin-dependent degradation, but the degron core, amino acids 82-94, doesn’t. In lanes 1-4, ectopic PAF53 was visualized with the anti-PAF53 antibody. In lanes 5 and 6, the AID-PAF53 was FLAG tagged and visualized with anti-FLAG antibody. D. Amino acids 68-111 program PAF53 for auxin-dependent degradation if placed on either the N-or C-termini of the protein. The TIR1 used in these experiments did not contain a nuclear localization signal. The analyses presented were carried out after three hours of auxin treatment (500 μM of IAA (+ auxin) or water (– auxin). Western blots for PAF53 were carried out as described (48).

Based on these observations, we established cell lines that constitutively expressed TIR1. However, we found that these cell lines did not degrade AID-tagged PAF53 as rapidly as predicted (data not shown) from the transient transfection experiments. We considered two models. Either we did not have the same levels of TIR1 expression in the stable cell lines as in the transiently transfected cell lines, or the TIR1 was not localizing to the nucleus/nucleolus. We examined the sequence of the TIR1 we were using and found it did not contain a nuclear localization signal (NLS). An NLS was added to the N-terminus of TIR1 and cells that constitutively expressed NLS-TIR1 were selected with hygromycin. One clone, 569, that demonstrated the rapid degradation of transiently transfected PAF53-AID was chosen for further cloning experiments. Clone 569 was then subject to CRISPR driven homology directed repair to introduce the AID onto the C-terminus of the endogenous PAF53 genes using the strategy summarized in Figure 2A. After selection and cloning by limiting dilution, the clones were screened by western blots for expression of the chimeric PAF53 to select for cells homozygous for AID-PAF53 expression (Figure 2B).The two forms of PAF53 seen in lane 1 is the result of tagging only one allele and lane 2 presents the results of tagging both alleles. Further, recombination was confirmed by PCR for the recombinant insert (Figure 2C). The PCR products were cloned and four clones were sequenced. The sequences were identical, and did not demonstrate errors in recombination (data not shown).

**Figure 2.**
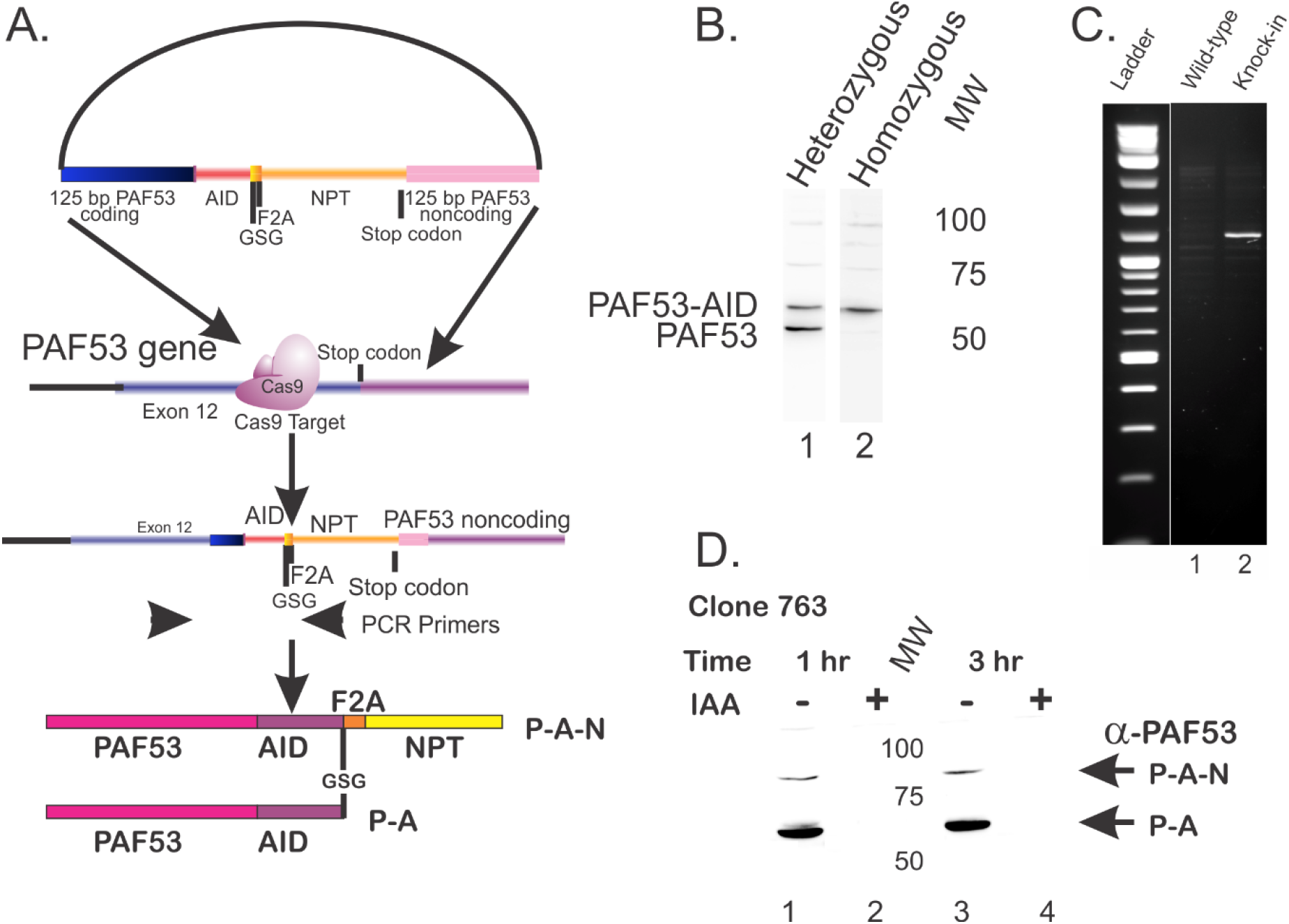
Targeting PAF53 with an AID. A) Design of the oligonucleotide used to recombine the AID sequence into the endogenous PAF53 gene. The oligonucleotide contained 125 bp of exon 12 of the human PAF53 gene in frame with aa 68-111 of IAA17. This was followed, in frame, by GSG, an F2A sequence and DNA coding for neomycin phosphotransferase (NPTII) and a stop codon and 125 bp of 3’ noncoding sequence from the PAF53 gene. The oligonucleotide was cloned into pUC57Kan, and intact plasmid was co-transfected along with pLentiCRISPRv2 as described in Materials and Methods. Stable recombinants were selected with G418 and subject to dilution cloning. (B) Examples of monoallelic recombination (lane 1) and biallelic recombination (lane 2) as determined by western analysis with anti-PAF53 antibody. C) PCR analysis of homozygous recombinants using the PCR primers indicated in Panel A. D) Treatment with 500 *μ*M IAA results in the degradation of PAF53-AID in one hour. The cells of Clone 763 demonstrated two proteins that reacted with anti-PAF53 antibody. The major band, migrating at ~ 58 kDa, was at the molecular mass predicted for the AID-tagged PAF53 (P-A). The upper band migrated with a mass predicted if it were the complete PAF53-AID-P2A-NPT chimeric protein (P-A-N), ~ 87 Kda,

Clones that only expressed the AID-tagged PAF53 were selected, expanded and stored. We found that the anti-PAF53 antibody recognized two bands in some of the clones (Figure 2D). The major band, migrating at ~ 58 kDa, was at the molecular mass predicted for the AID-tagged PAF53 (P-A). The upper band migrated with a mass predicted if it were the complete PAF53-AID-F2A-NPT chimeric protein (P-A-N), ~ 87 Kda, due to incomplete F2A-dependent cleavage. When IAA was added to these cells, we observed a rapid degradation, within one hour, of both bands that reacted with anti-PAF53 antibodies (Figure 2D, lanes 2 and 4). This confirmed that they were the bands predicted and also demonstrated that the presence of an AID in the middle of a protein could target the protein for degradation.

As introduced, the role(s) of yeast A49 in rDNA transcription in rDNA transcription are not clear. When mammalian cells were screened for genes “required for proliferation and survival”, Wang *et al*. reported that the PolR1 E (PAF53) gene was essential. This result was confirmed by Bertomeu *et al*. (16) and others (50). Thus, we sought to determine the specific effect of knocking down PAF53 using the AID as both physiological and biochemical results should be apparent very rapidly.

As shown in Figure 3A, treatment of clone 763 that expressed both TIR1 and AID-tagged PAF53 for three hours with IAA resulted in the inhibition of rDNA transcription as demonstrated by metabolic labeling of total cell RNA. This experiment has been reproduced at least three times and we observed at least 99% inhibition of synthesis of the 47S pre-rRNA in all experiments. The inhibition of rDNA transcription can induce a phenomenon referred to as “nucleolar stress” or “ribosome stress”. The cellular responses to this stress can include cell cycle arrest or cell death (77–79,89–94). When clone 763 cells were exposed to IAA, they appeared to complete one round of division and then arrest (Figure 3B). The cell cycle arrest was stable for up to nine days with no apparent increase in cell death. To confirm that the cell-cycle arrest was due to the knock-down of PAF53, we sought to determine if the ectopic expression of the protein would rescue the arrest. Clone 763 cells were transfected with pCDNA3.1 driving the expression of mouse PAF53 and a vector driving the expression of eGFP two days before the cells were treated with IAA. Control cells were transfected with an empty vector and the vector driving eGFP. Subsequently, the number of fluorescent cells was measured in each of the three groups. As shown in Figure 3, expression of ectopic PAF53 rescued rDNA transcription (Figure 3A, lanes 4) and prevented cell cycle arrest (Figure 3B). Western blot analysis (Figure 3C) confirmed the expression of the mouse homolog and the lack of expression of the endogenous AID-PAF53 throughout the time course of the experiment.

**Figure 3.**
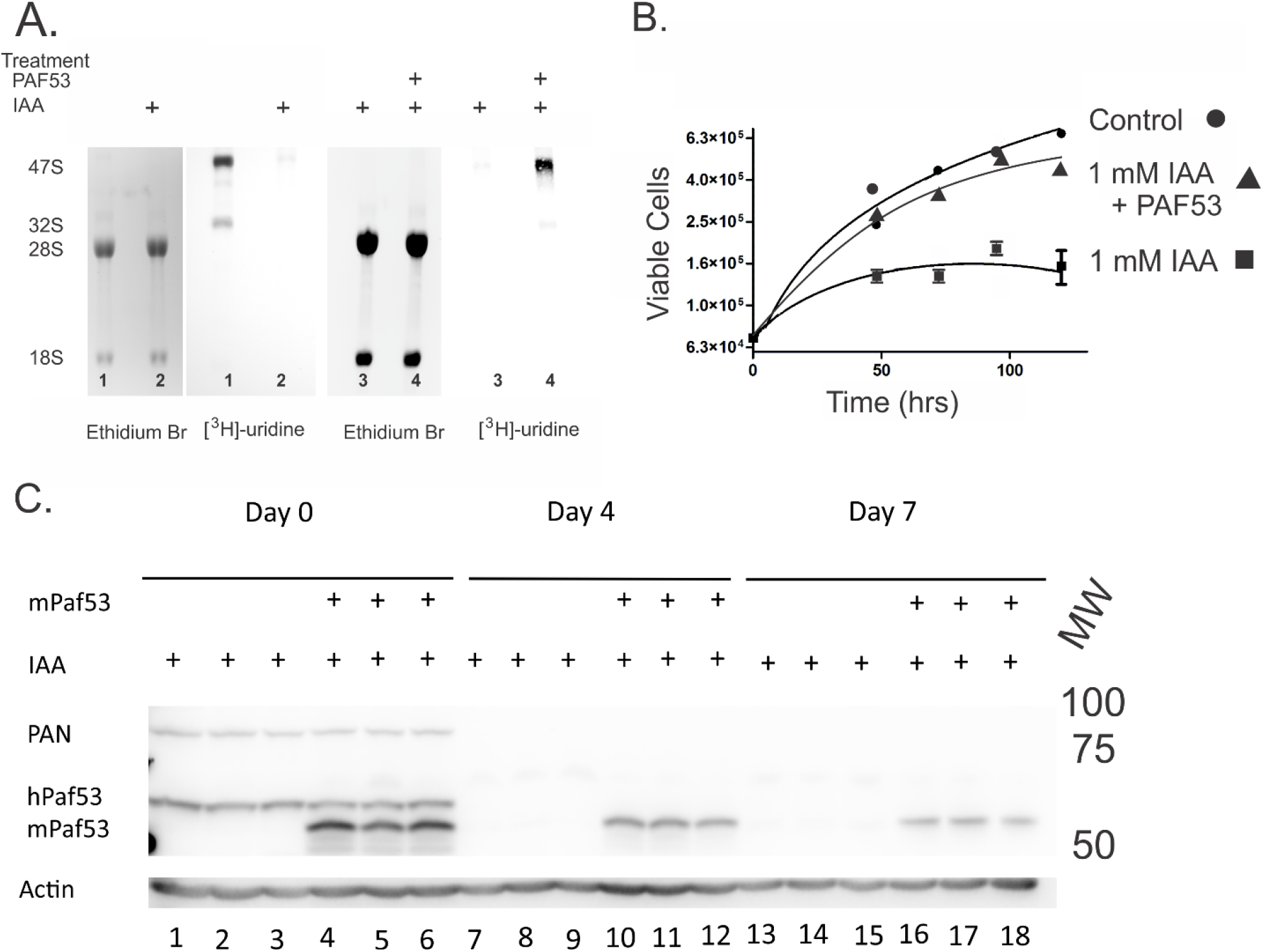
Depletion of the endogenous PAF53-AID results in (A) the arrest of cell proliferation and (B) the inhibition of rDNA transcription which can be rescued by the ectopic expression of PAF53. A) Cells cultured in 30 mm wells were treated with 500 *μ*M IAA at “0” time. All cells were transfected at −48 hr with either empty vector (control) or pcDNA3.1 expressing wild-type mouse PAF53 (+PAF53). Cells were then harvested at the times indicated and the number of cells determined as described (116). The growth curves were fitted with a polynomial. N=3, +/− Std. Dev. B) Metabolic labeling demonstrates the inhibition of pre-47S rRNA synthesis upon depletion of PAF53.Clone HEK763 that expressed both TIR1 and AID-tagged PAF53 were treated with 500 *μ*M IAA for three hours. Following treatment, [^3^H]-uridine was added to the media for 30 minutes and total RNA was isolated an analyzed as described in Materials and Methods (116). (C) The expression of ectopic mouse PAF53 is maintained over the seven day period of the experiments described in Panel A. Cells were treated with IAA as indicated and harvested immediately after the onset of treatment or at the times indicated. The cells were lysed in SDS-sample buffer and the expression of endogenous or ectopic PAF53 determined by western analysis with anti-PAF53 antibody.

While it was clear that clone 763 cells depleted of PAF53 did not demonstrate a net proliferation, this could be the product of cell cycle arrest or a steady-state relationship between cell division and cell death. In order to examine this question, we examined the cells for trypan blue dye exclusion. We did not observe a significant increase in trypan blue staining following treatment with IAA, indicating that we were not causing cell death (data not shown). The inhibition of rDNA transcription has been shown to cause cell cycle arrest at the M/G1 boundary. However, the effect of rapidly targeting rDNA transcription has not been previously studied. Previous studies have used Cre-dependent recombination to knockdown RRN3 and UBF (95,96), two components of the rDNA transcription apparatus (97). This process takes much longer than an hour and can cause apoptosis (96). As shown in Figure 4, twenty-four hours after the depletion of PAF53, there was no alteration in the distribution of cells through the cell cycle. However, after two days, we observed a decrease in the percentage of the cells in G1 and an increase in the percent of cells in S (Figure 4B). The increase in the S/G1 ratio was significant after 48 hours (Figure 4C). Interestingly, we did not observe a significant accumulation of cells in either the G1 or G2 checkpoints. This is in contrast to the patterns observed when RRN3 and UBF were knocked down (95,96). In those instances the investigators observed accumulation of the cells in either G1 (RRN3) or G2 (UBF) concomitant with a minor inhibition of the distribution of cells in S or its abrogation.

**Figure 4.**
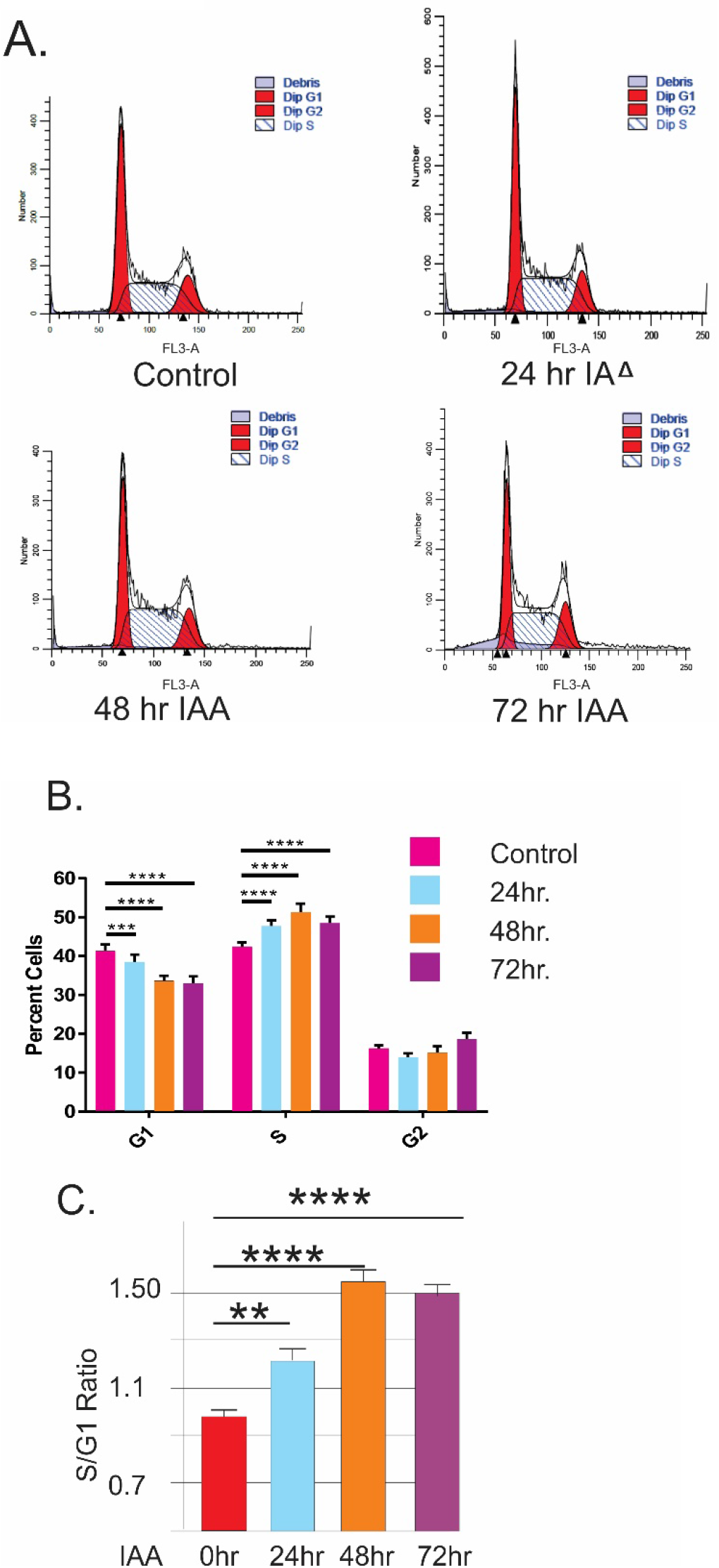
Depletion of PAF53 causes an increase in the S/G1 ratio of cell cycle distribution. (A) FACS analysis of cells at the indicated times post exposure to IAA. (B) Quantitation of the distribution of cells in G_1_, S or G_2_ following IAA treatment. Three independent repeats of the analyses presented in Panel A were each carried out in triplicate. The data were analyzed by one-way ANOVA. (C) The ratio of cells in S and G1 was calculated for the data presented in Panel B. Significance was determined by a one-way ANOVA. *= 0.05-0.01; **= 0.01 −0.001; ***= 0.001 −0.0001;****= <0.0001.

As shown in Figure 5, yeast A49 can be considered to consist of three domains an N-terminal dimerization domain that mediates the interaction with A34, a C-terminal tandem winged-helix (t-WH) that has DNA-binding capacity and a linker domain between the two [(88) and SWISSMODEL, Q9GZS1, see also PDB entry 3NF1]. Both SWISS MODEL and I-TASSER predict structures for mouse and human PAF53 similar to that of yeast A49 despite <25% sequence identity (98–102). We have previously demonstrated that the N-terminal domains of mouse PAF53 and PAF49 are required for dimerization, and are predicted to contain the triple barrel *β*-structure found in the A49/A34 heterodimer and in TFIIF (42,44,88,103). Interestingly, I-Tasser and SWISSMODEL (Q9GZS1) predict similar structures for the C-terminal domains of the yeast (NP_014151.1) and mammalian homologues (human NP_071935.1 or AAH14331.1) despite their being only 22% identical. While it has been demonstrated that the t-WH of yeast A49 has DNA-binding activity, the same has not been reported for the predicted t-WH of the mammalian homologues. As shown in Figure 6A, we determined that human PAF53 was capable of binding to DNA. Moreover, we found that the N-terminal dimerization domain was not required for DNA-binding (Figure 6B. aa109-435; lanes 2 and 3). The t-WH domain, amino acids 216-435, also bound DNA (lanes 4 and 5). However, when the N-terminal WH was deleted, the remainder lost DNA-binding activity (lanes 6 and 7). Thus, both the yeast and mammalian homologues contain N-terminal dimerization domains and C-terminal DNA-binding domains.

**Figure 5.**
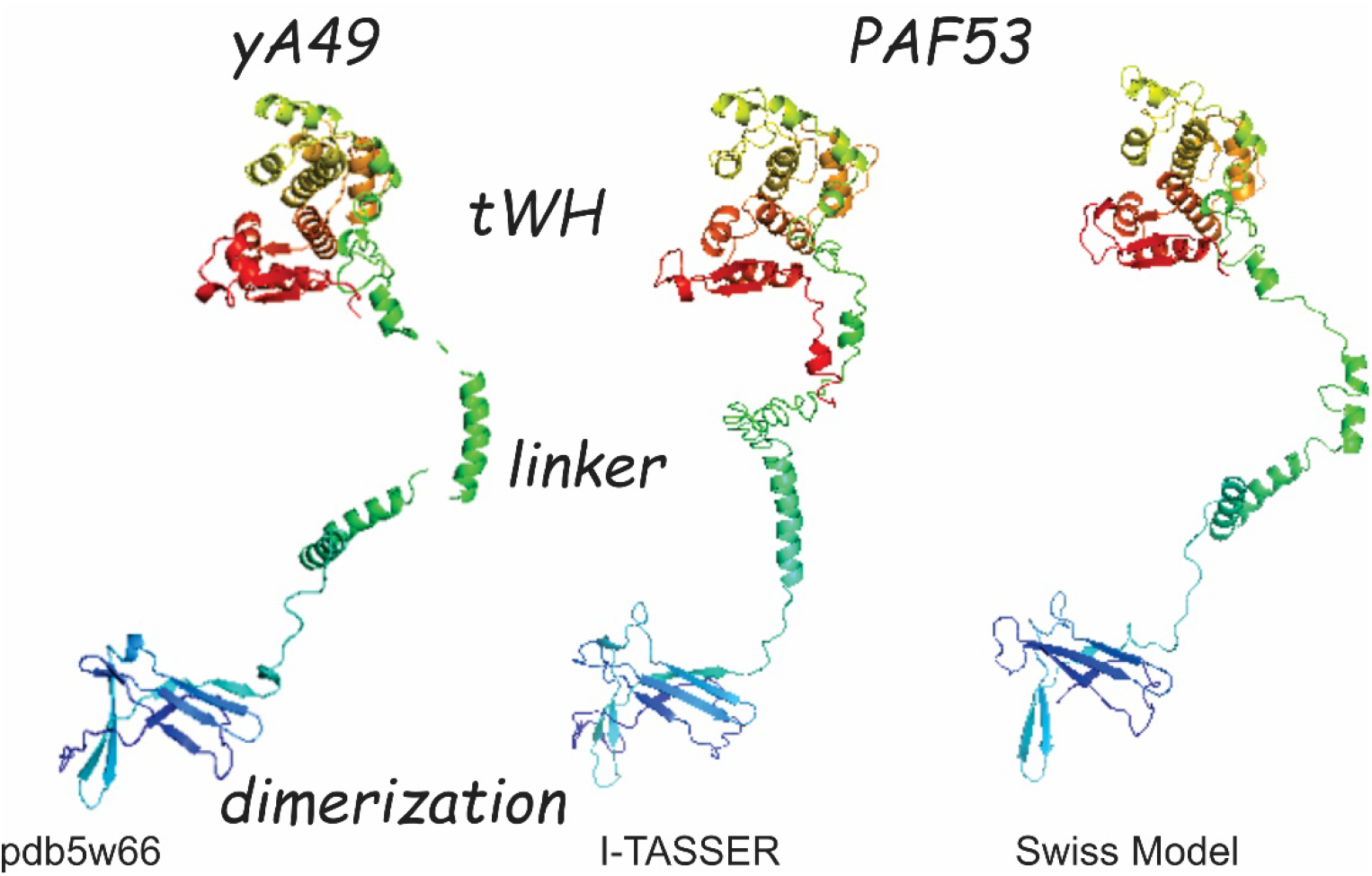
Domains of Yeast A49 and Human PAF53. The structure of yeast A49 was adapted from pdb5w66. The structures of human PAF53 were generated by SWISS MODEL (98,99) and I-TASSER (101,102,109) as indicated. The dimerization and tWH domains for yeast A49 are indicated. Similar structures are apparent in the predicted structures of human PAF53. The domains of yeast A49 are adapted from Geiger *et al*. (42)

**Figure 6.**
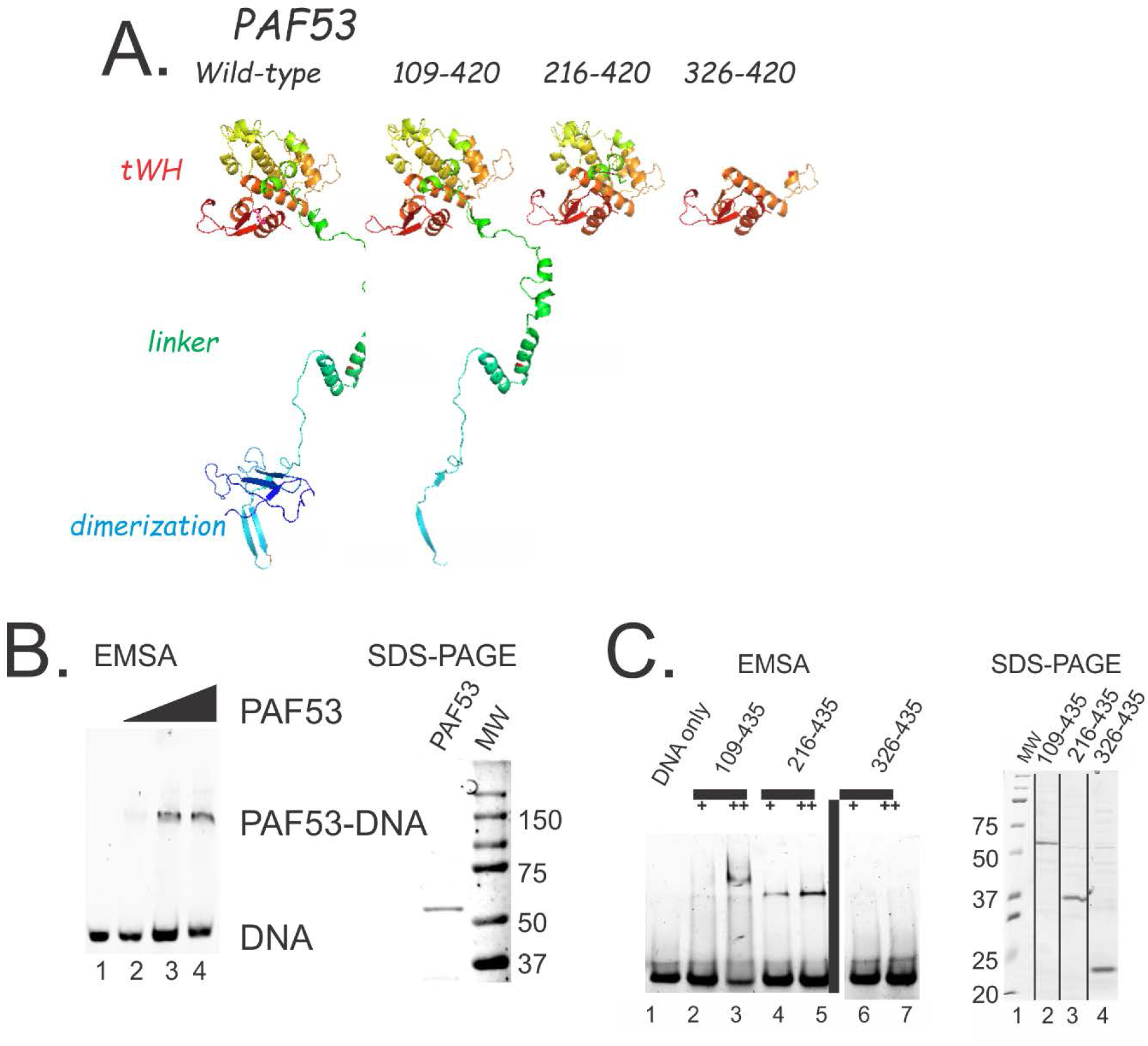
The putative tWH structure of mouse PAF53 has DNA-binding activity. (A) SWISS-MODEL predictions for the structures of the PAF53 constructs used in these experiments. The predicted structure generated by SWISS-MODEL terminates at amino acid 420. (B) Purified, full length mouse PAF53 has DNA-binding activity. DNA-binding assays with increasing amounts of PAF53 are presented in lanes 2-4 of the right hand figure in panel B. A coomassie stained gel of the protein used is presented in the left hand figure in panel A. (C) Deletion mutagenesis of PAF53 demonstrates that the PAF53 lacking the N-terminal dimerization domain has DNA-binding activity and that mutagenesis of the tWH inhibits DNA-binding. The indicated mutants were used in the band-shift assays in two different concentrations (+, ++) in the assays. The amounts of the proteins used in the EMSA were normalized to the coomassie stained bands. The conditions for the EMSA were as described in Materials and Methods.

We then sought to determine which, if any, of the domains of the protein were essential for cell cycle progression. Two days prior to treatment with IAA, cells were transfected with vectors expressing various deletion mutants of mouse PAF53 (Figure 7A). We found that a construct lacking the N-terminal dimerization domain (PAF53 aa109-435) failed to restore cell division (Figure 7D). The N-terminal dimerization domain (PAF53 aa1-160) itself did not rescue cell division either (Figure 7C). Surprisingly, we observed partial rescue with a construct that did not contain the C-terminal tWH, but did contain most of the dimerization domain and the linker (PAF53aa1-222) (Figure 7B). These three constructs would appear to be properly folded, as deletion of the dimerization domain did not inhibit DNA-binding by the t-WH (Figure 6B lanes 4 and 5) and deletion of the t-WH did not inhibit dimerization (88). Further, all three constructs were expressed for the duration of the experiment (Figures 7E and F). Thus, the linker region between the dimerization and tWH domains appears to be a functional domain.

**Figure 7.**
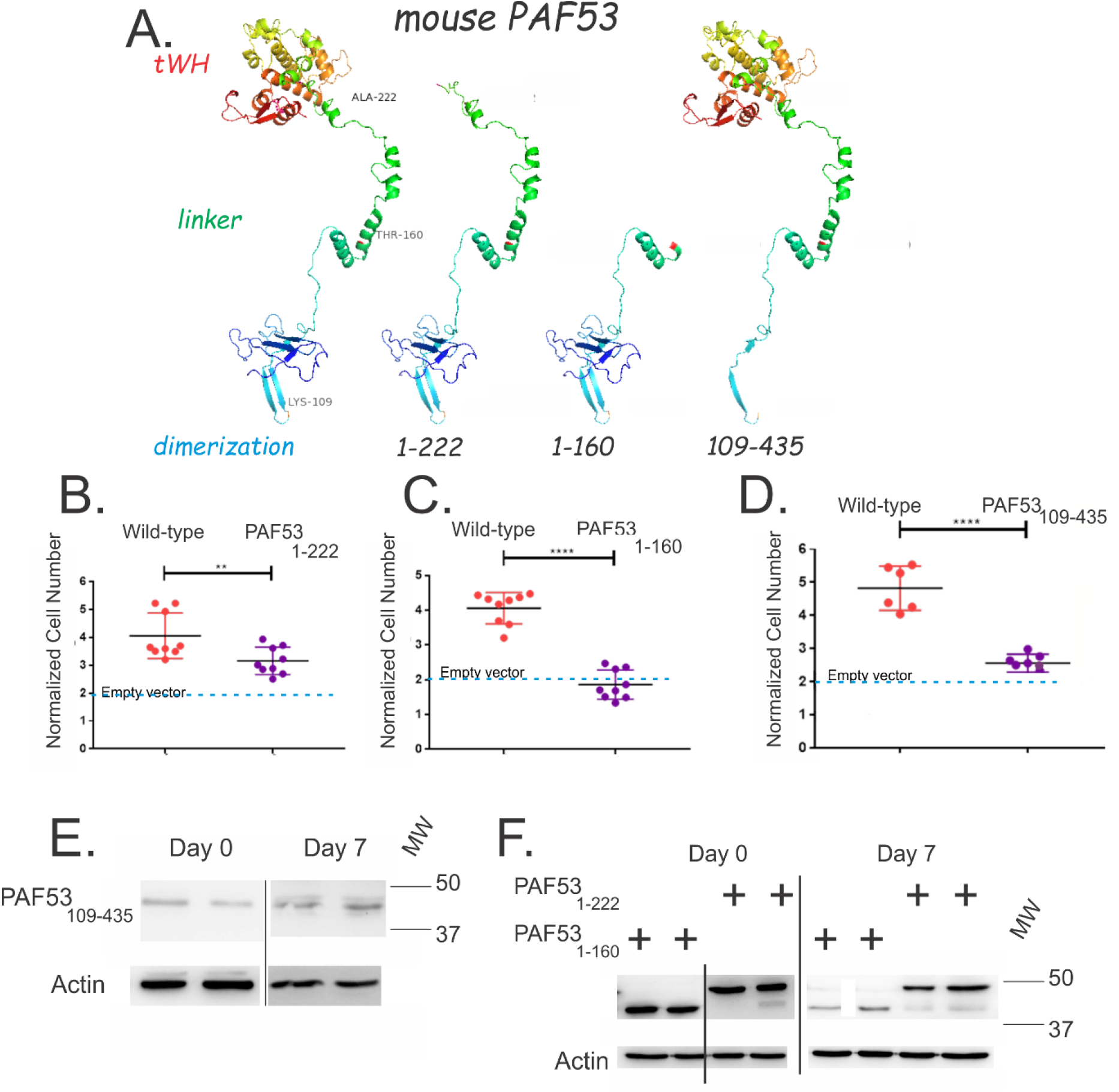
PAF53 function requires all three domains of the protein. Forty-eight hours prior to treatment with IAA, cells were ctransfected with vectors expressing eGFP and either wild-type PAF53 or the indicated deletion mutants of mouse PAF53. Cells were then treated with IAA (500 *μ*M) to deplete endogenous PAF53. Five days after IAA, cells were harvested and counted. The growth of the cells transfected with mutant forms of PAF53 is presented relative to the growth of cells transfected with wild-type PAF53 in matched experiments. (A) PAF53 constructs used in the rescue assays. The predicted structure generated by SWISS-Model terminates at amino acid 420. (B) In comparison to wild-type PAF53, PAF53 1-222 was approximately 80% as active in rescuing cell division. (C) Further deletion of the linker domain inhibits rescue. PAF53_1-160_ did not rescue cell division. (D) The N-terminal dimerization domain is required to rescue cell division. Deletion of the N-terminal dimerization domain, as in PAF53_109-435_ inhibits the ability of PAF53 to rescue cell proliferation. The yellow bars indicate the cell number that was observed when the cells were treated with IAA and transfected with an empty vector. (E and F) Western blots for PAF53 demonstrate that all three of the deletion mutants tested were expressed for the duration of the experiment. Significance was determined by a two-tailed t-test.

When we examined the structure of the linker, we noted that both mammalian PAF53 and yeast A49 contain a helix-turn-helix motif (HTH). It has been reported that the linker spans the cleft in Pol I (31,104). We hypothesized that the HTH might come in contact with the template and that the linker might participate in transcription due to DNA-binding activity. When we expressed His-PAF53 aa1-222 and tested it in a DNA-binding assay, we found that the purified, recombinant protein had DNA-binding activity (Figure 8). To examine the role of the putative HTH in DNA binding, we mutated each of the helices individually (Figure 9, panels A and B) and tested equal amounts of the recombinant proteins for DNA binding activity. As shown in figure 9, panel C, mutation of either of the helices significantly inhibited DNA binding. Thus, mammalian PAF53 has two domains with DNA-binding activity, the tWH and the helix-turn-helix of the “linker”. Subsequently, we carried out rescue experiments using PAF53 constructs that contained similar mutations of the HTH. As shown in Figure 10, cells expressing PAF53 with the HTH mutated failed to divide following IAA treatment.

**Figure 8.**
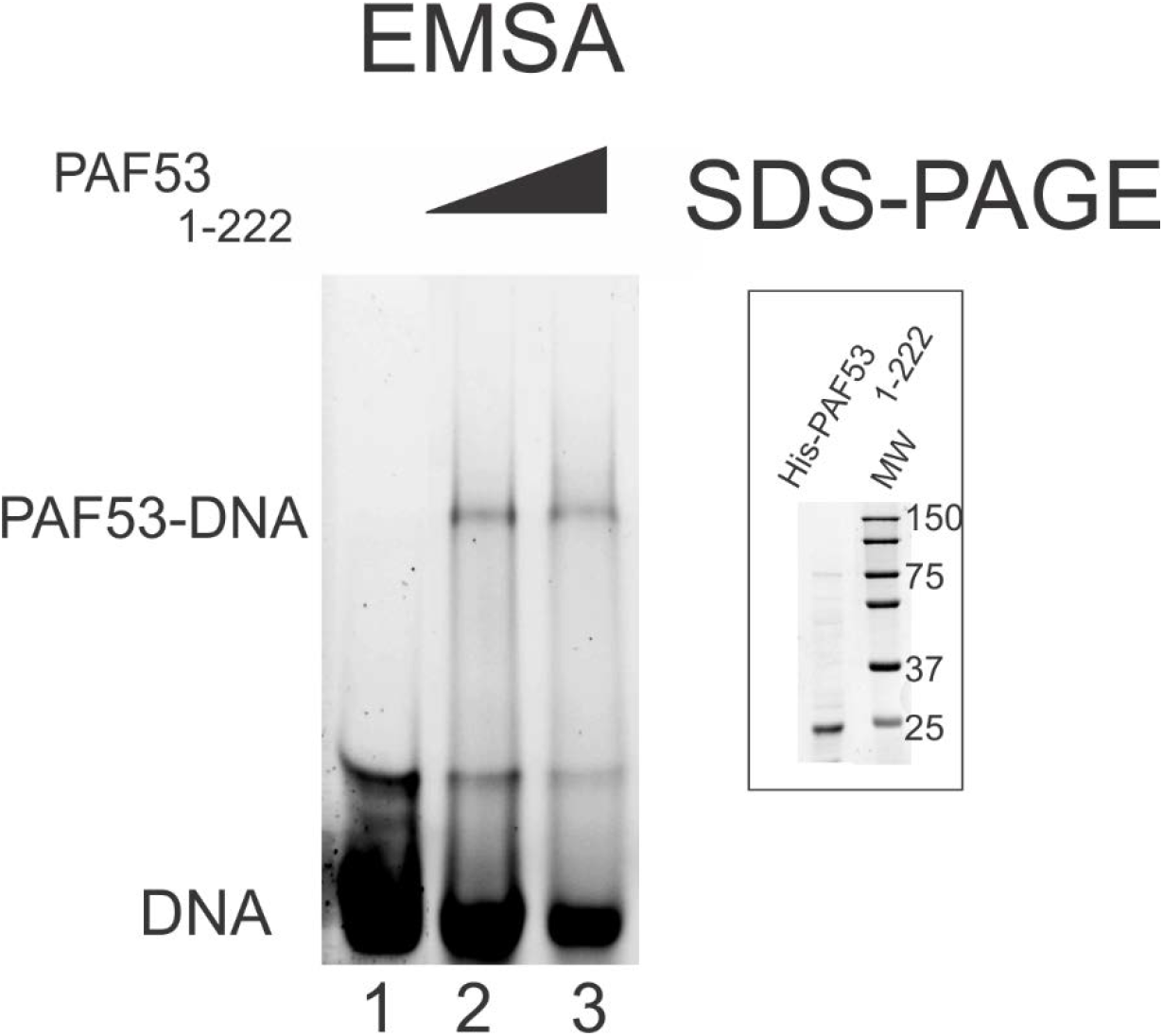
Amino acids 1-222 of PAF53 have DNA-binding activity. His-PAF531-222, lacking the C-terminal tWH domain, was expressed in BL21, purified by IMAC, analyzed by SDS-PAGE (inset) and assayed for DNA binding activity using the same fragment as in Figure 6.

**Figure 9.**
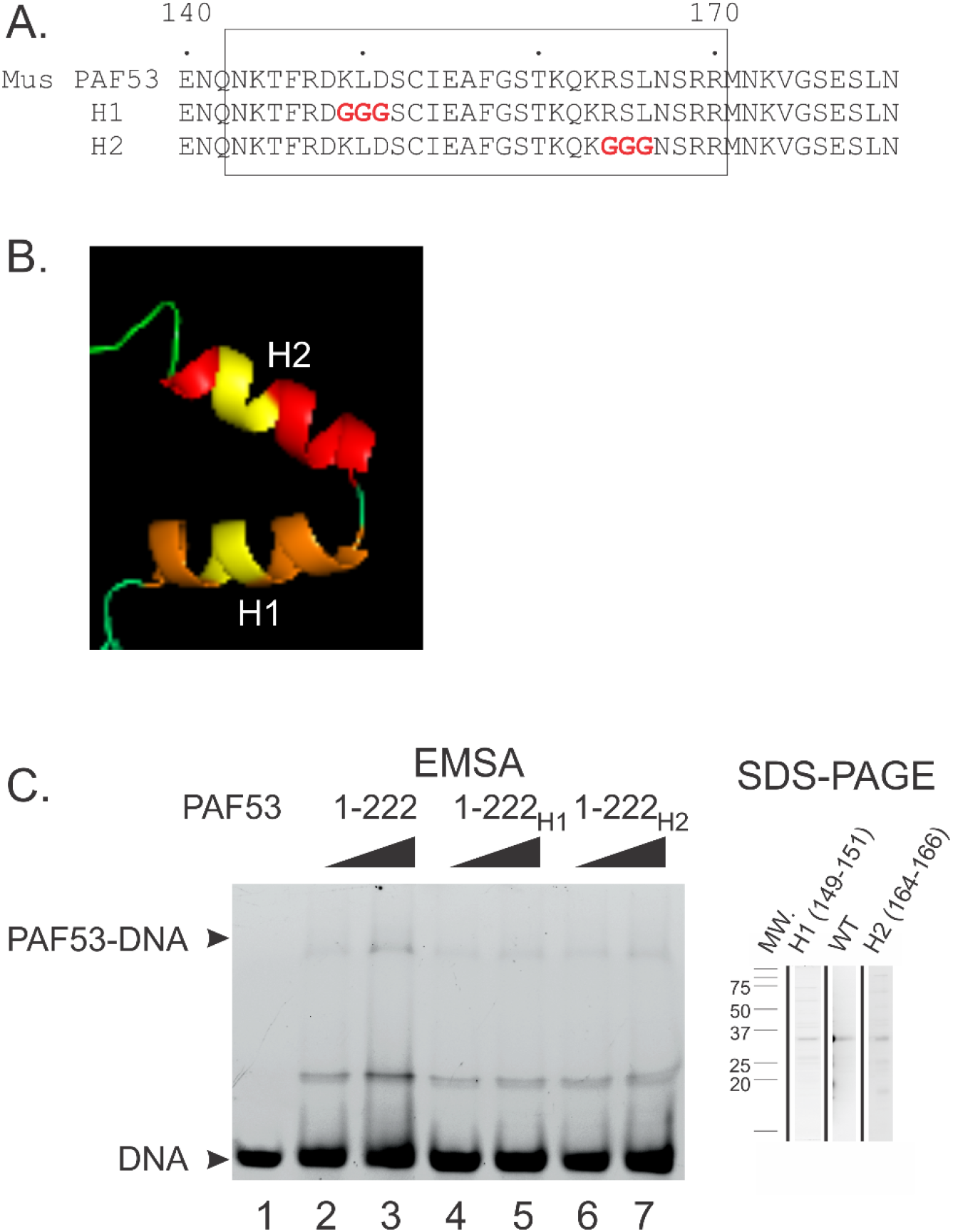
The helix-turn-helix domain of PAF53 is responsible for DNA-binding activity. A. Sequences of wild-type and mutant forms of PAF53 1-222. I-tasser prediction of the structure of PAF53 amino acids 140-180 demonstrating (in yellow) the glycine substitution in the two predicted helices. C. Mutation of amino acids 149-151 (H1) or 164-166 (H2) (glycine substitution) reduces DNA-binding activity demonstrated by amino acids 1-222 of PAF53. D. Simultaneous mutation of H1 and H2 inhibits DNA-binding by PAF53_1-222_ and reduces DNA-binding by full length PAF53. The various mutants were of His-PAF531-222, lacking the C-terminal tWH domain, were expressed in Rosetta DE3 (Novagen), purified by IMAC, analyzed by SDS-PAGE (insert) and assayed for DNA binding activity using the same fragment as in Figure 6.

**Figure 10.**
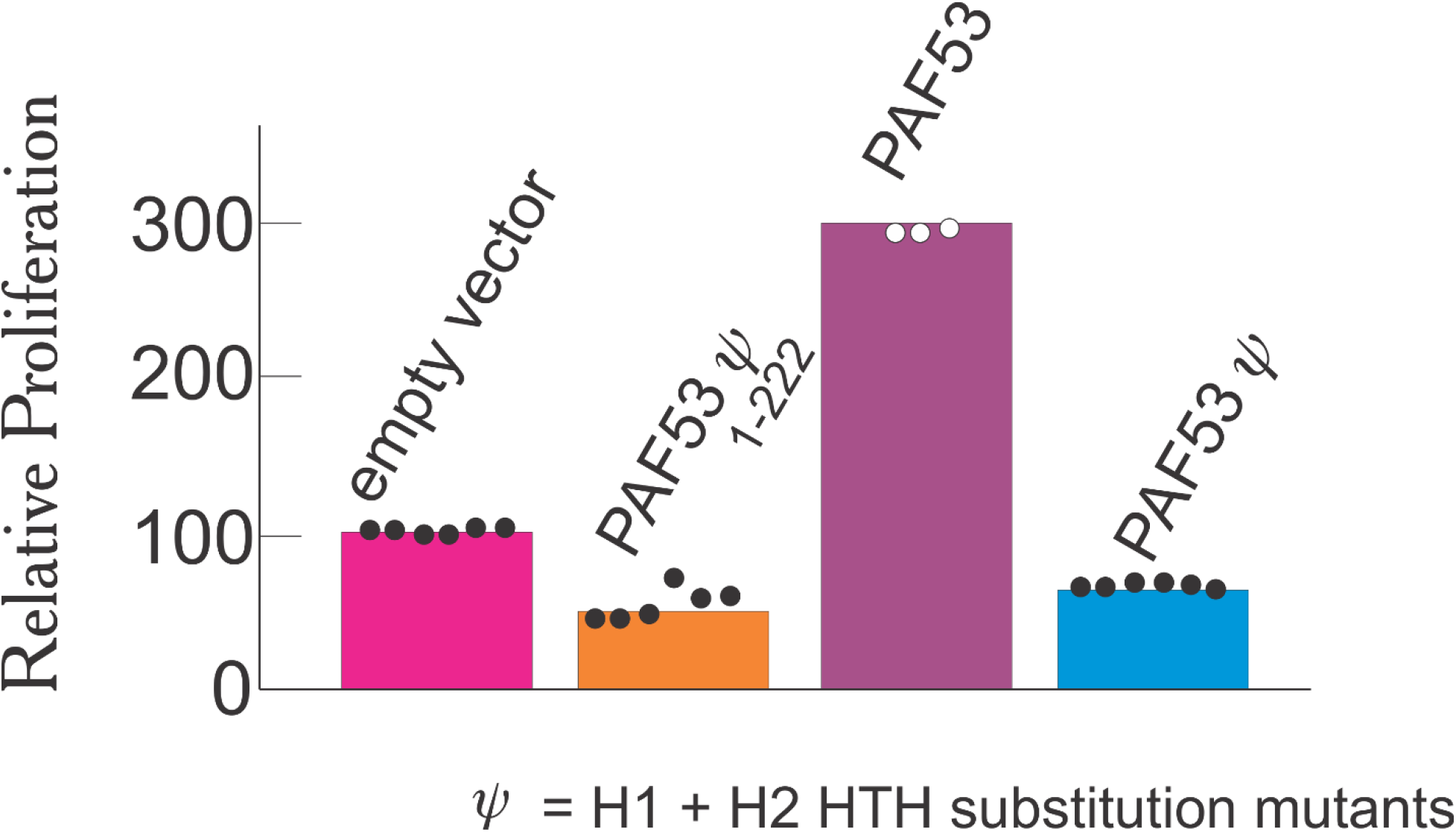
PAF53 function requires the linker HTH to function. Forty-eight hours prior to treatment with IAA, cells were cotransfected with vectors expressing eGFP and either wild-type PAF53 or the indicated HTH mutants of mouse PAF53 or mouse PAF531-222. Cells were then treated with IAA (500 μM) to deplete PAF53. Five days after IAA, cells were harvested and GFP+ cells counted. The growth of the cells transfected with the indicated forms of PAF53 is presented relative to the growth of cells transfected with empty vector in matched experiments (n≥3).

## Discussion

We have established that the TIR1, auxin-dependent degron system can be used to rapidly deplete cells of nucleolar proteins. Our data demonstrate the rapid (<1hr) auxin-dependent degradation of a nucleolar protein, PAF53 and the feasibility of using this approach to study the function of those proteins in mammalian cells following depletion or replacement with mutant forms of the protein. In our initial investigations of the feasibility of this system we found that we had to optimize the system in order to apply it to nucleolar proteins.

Several laboratories have used the TIR1 from *Oryza sativa* (osTIR1) to target nuclear or cytoplasmic proteins in mammalian cells as it is more stable at 37C than the TIR1 from *Arabidopsis thaliana* (83). This was the form of TIR1 we started with. When TIR1 was expressed in HEK293 cells, we did observe depletion of AID-PAF53. However, the cells had to be exposed to auxin for at least two hours before we saw nearly complete depletion. When we analyzed the sequence of the osTIR1 we noted that it did not contain a nuclear localization sequence. To target osTIR1 to the nucleus, we added a monopartite NLS (PAAKRVKLD) to the N-terminus using PCR mutagenesis. Western analysis confirmed that NLS-osTIR1 was active; it degraded endogenous AID-tagged PAF53 within one hour.

Next, we considered the design and placement of the degron. Initial studies on the use of the degron focused on placing it on the N-terminus of the targeted protein, as it works in plants. In agreement with Nishimura and colleagues we found that the degron also functions when placed on the C-terminus (80). Thus, the degron works when placed on either terminus. Interestingly, it appears that the degron also works when placed internally. The construct we used to target PAF53 for degradation results in the expression of a protein that contains PAF53-AID linked through a F2A or *cis-* acting hydrolase element, CHYSEL site (105–107), to neomycin phospho-transferase (P-A-N). Not all of the precursor molecules were cleaved and there was residual P-A-N apparent on the western blot. When the cells were treated with IAA the P-A-N was degraded suggesting that the AID functions internally.

The next question we addressed was the choice of degron. The full length degron of 229 amino acids as found in auxin-responsive protein IAA17 (IAA17, indole-3-acetic acid inducible 17) of *Arabidopsis thaliana*, has been demonstrated to target proteins for degradation (80–82), and we confirmed its use in several cell lines (data not shown). However, we thought it was possible that the size of this degron might interfere with protein function in future experiments. Several laboratories have reported that the 13 amino acid conserved core of domain II of the degron is sufficient to support auxin-dependent degradation [see ref. (82)]. However, when we coupled that sequence to PAF53 we did not observe rapid degradation of the protein in mammalian cells. There was a third option. An intermediate sized fragment of domain II of IAA17 has been shown to function as a degron in yeast (108). When we coupled aa 68-111 of IAA17 to the C-terminus of PAF53 we found that the tagged protein was degraded within an hour. This is the degron that we have used for the studies presented in this manuscript.

The yeast A34 and A49, and their mammalian homologs, PAF49 and PAF53, are structurally similar. Figure 5 compares the structures of yeast A49 and the structure predicted for human PAF53 by I-TASSER (101,102,109). The three domains, dimerization, linker and tandem winged helix (tWH), of A49 are indicated (88).

We have previously mapped the regions in PAF53 and PAF49 that mediate heterodimerization. The interaction of PAF53 with PAF49 requires the first 160 amino acids of PAF53 (88) and the amino acids 41-86 of PAF49. *In silico* analysis of the heterodimerization domains found that their amino acid sequences were 21% identical to the yeast homologs, but structurally similar to the heterodimerization domains of A34 and A49 (88). We have previously reported that when PAF49, the mammalian homolog of A34, is expressed ectopically it binds to Pol I in the absence of ectopic PAF53. Similarly, when PAF53 was expressed ectopically, it demonstrated binding to Pol I (88). Our observation that a form of PAF53 which lacks the heterodimerization domain (PAF53 aa109-435) does not rescue cell growth is strongly supportive of a model in which a significant portion of the interaction of PAF53 with Pol I is dependent upon the dimerization domain and perhaps binding to PAF49. That is, the interaction of mammalian PAF53 with PAF49 is necessary for PAF53 function.

The tWH domain of A49 has been demonstrated to be a DNA-binding domain, and models of rDNA transcription suggest that the tWH of A49 may have multiple roles in rDNA transcription including the stabilization of the open complex by binding upstream DNA (42,110). We have found that the putative tWH of mammalian PAF53 has DNA binding activity and that the C-terminus of PAF53 is required for full activity. this suggests an essential role for the tWH in mammalian cells. Engel *et al*. suggest that the tWH domain may move with upstream DNA facilitating the transition from initially transcribing complex to elongating complex and stabilizing that structure(111). This would be consistent with the finding that the tWH domain enhanced processivity *in vitro* (42).

As shown in Figure 5C, we found that PAF53_1-222_ was nearly eighty percent as effective as intact PAF53 in its ability to support cell division. In contrast, a mutant that lacks a significant fraction of the linker (aa1-160) was essentially inactive in this assay. When combined with the finding that the dimerization domain itself was necessary to fully rescue cell growth, these results would argue for an essential role for the linker.

Experiments on yeast A49 had not predicted a role for the “linker”. The observation that *in silico* analysis of the linker of mammalian PAF53 predicts a structure similar to that observed for the yeast argues for a conserved function. The predicted structure for mammalian PAF53 contains a helix-turn-helix domain (HTH), a potential DNA-binding domain, that lies very close to the DNA just downstream of where the bubble would form in the closed complex (PDB 5w66). Further, the construct PAF53_1-160_ clips the c-terminal helix within this HTH. We then found that this region has DNA-binding activity, and that mutagenesis of the helices, alone or in combination, inhibited DNA-binding by PAF53_1-222_, Interestingly, the structure of the linker has not always been clearly discerned in structures of Pol I (104), possibly due to dynamics in its interactions with core Pol I or the template. It has been proposed that the linker spans the active site cleft and may in fact play a role in the narrowing of the cleft in the process of proceeding from initiation to escape (104,112). We are determining the role of the HTH in transcription, *e.g*. the transition from initiation to elongation.

Our results demonstrate that PAF53 is necessary for transcription by Pol I and mitotic cell growth. These results support the model that rDNA transcription is required for cell proliferation (1,113–115). This study has shown that all three domains of PAF53 are necessary for necessary for mitotic cell growth. The role(s) each of these domains play during rDNA transcription will be the subject of future studies. Most particularly we found a requirement for both the linker and the heterodimerization domains of PAF53 (88). This latter finding suggests that PAF53 function may be dependent upon the interaction with PAF49. As this would be different from what is observed in *S. cerevisiae* it bears further scrutiny. Previously, our lab has demonstrated that inhibiting rRNA synthesis in cancer cells causes cell death while normal cells arrest (116), and ribosome biogenesis has become a target for cancer chemotherapy (14,117–119). Understanding the role PAF53 plays in Pol I transcription could provide novel drug targets that could be utilized in cancer therapy.

## Experimental Procedures

### Cell Culture, Transfection, Selection and Analysis

HEK293 cells (ATCC) were cultured as recommended in DMEM containing 10% FBS and Invitrogen Antibiotic-Antimycotic. Cells were routinely passed at 1:4 dilutions every third day and were not used after the 18^th^ passage. Transfection of 60% confluent HEK293 cells was carried out as described (49,88) using PEI (120). After eight hours, the medium was changed. When selecting for stable transfections, the selection antibiotic was added 48 hr. post transfection. The sequences encoding NLS-TIR1 were cloned in pcDNA3.1/Hygro(+), and selection for expression of NLS-TIR1 required hygromycin (100 *μ*g/ml). When the goal was to recombine into the PAF53 gene, the cells were given puromycin 24 hours following transfection. Forty-eight hours later, the medium was changed to medium free of puromycin, but containing G418 (500 *μ*g/ml) to select for recombinants. In order to determine if specific constructs rescued the ability of cells to divide, the cells were cotransfected with a vector expressing the PAF53 construct to be tested and a vector expressing eGFP to mark the transfected cells. Two days later, 500*μ*M indole acetic acid (IAA) was added to the culture medium. IAA (Abcam) was made fresh by dissolving in water immediately before use. After the indicated time, the number of green fluorescent cells in the population was determined by FACS analysis. At least 10,000 cells were counted for each analysis, which was carried out in triplicate for each time point of an experiment. Cell counts, trypan blue exclusion and cell cycle analysis was carried out as described **(116,121)**. Metabolic labeling with [^3^H]-uridine was done as described previously (116). Briefly, clone 763 cells were transfected with empty vector or vector expressing mouse PAF53. 48 hours later, the cells were treated with IAA or vehicle. Three hours later, [^3^H]-uridine was added to the medium. After 30 minutes, the cells were harvested and RNA was isolated. The isolated RNA was fractionated by agarose gel electrophoresis, stained with ethidium bromide (Ethidium Br), photographed, transferred to PVDF. The filters were dried and impregnated with Enhance (Perkin Elmer) and subject to fluorography as described previously (116).

### Constructs

The wild-type and deletion clones of mouse PAF53 used in these experiments were described previously (49,88) and is NCBI Reference Sequence: NP_073722.1. In these clones, PCR was used to insert a FLAG-tag onto the N-terminus of wild-type mouse PAF53 cloned in pCDNA3.1 downstream of the CMV promoter. The deletion clones, 1-160 and 1-220 were cloned in pEBG-GST (Addgene). The vector expressing eGFP was described previously (49). All constructs were confirmed by sequencing. The vector expressing AID-H2B-YFP was obtained from Addgene.

### gRNA Design

We used CHOP**CHOP** (http://chopchop.cbu.uib.no/) as described (122) to design gRNA. Bsmb1 linkers were added to two gRNA targeting exon 12 of human PAF53, TAATCTTCCTCCGCTTTGCC<u>AGG</u> and TAGACGCATGCTTTCCAGAC<u>AGG</u> (the PAM is underlined), and the oligos were then annealed following a standard protocol and ligated into the vector (123,124), plentiCRISPR v2 (Addgene, (125)). The oligonucleotides were supplied by MilliporeSigma. The constructs were confirmed by sequencing. The use of plentiCRISPR v2, which is constructed around a 3rd generation lentiviral backbone, allows for the simultaneous infection/transfection of the vector for the expression of Cas9 and gRNA and for selection for puromycin resistance.

### Homologous Recombination

An oligonucleotide was designed that contained 125 nucleotides of PAF53 coding sequence in frame with DNA that codes for amino acids 68-111 of *A. thaliana* IAA17. This was followed by GSG and the sequence coding for a F2A self-cleaving peptide (105,106,126) (VKQTLNFDLLKLAGDVESNPGP) which was upstream of the sequence coding for neomycin phosphotransferase, a stop codon and the next 125 nucleotides of 3’ noncoding PAF53 exon (Figure 2). The oligonucleotide was synthesized by Genewiz and cloned into pUC57Kan. Cells were transfected with plentiCRISPRv2 coding for the gRNAs that target exon 12 along with the plasmid containing the oligonucleotide for recombination. Twenty-four hours following transfection, puromycin was added to the culture medium (6 *μ*g/ml). After another 48 hr., the medium was changed to medium containing G418 (500 *μ*g/ml) to select for recombination. After 72 hr., the surviving cells were subject to cloning by limiting dilution and the clones were expanded to 60 mm dishes. After cloning, the success of recombination was confirmed by PCR, as described previously, using the forward and reverse primers, 5’-CAGTCATGTTGAGGGGTCTCTCCAGTTCTTCTG-3’ and 5’-GCCCTTGAAGAACTCGTCAAGAAGGC - 3’ primers indicated in Figure 2 (50) and western blots for PAF53. The PCR products were cloned in PCR blunt II TOPPO (Invitrogen). Both strands of four separate clones were sequenced. Analysis demonstrated that the sequence of all four clones were identical.

### Western Blotting, Protein Purification and EMSA

Cell lysis and western blotting was carried out as described (127) using antibodies to PAF53 (48). The anti-PAF53 antibodies were either raised to recombinant PAF53 in our laboratory (48) or obtained from Proteintech Group and recognizes both human and mouse PAF53. The antibody from our laboratory has been validated previously (35,48), and the depletion of endogenous PAF53 results in a loss of immunoreactivity for both proteins. Recombinant proteins were expressed in either BL21 or Rosetta (Novagen) cells, purified as described previously using either His-, FLAG- or GST-tags (49,55,88) and analyzed by SDS-PAGE and Coomassie Brilliant Blue R staining. To quantitate the amount of protein used in the electrophoretic mobility shift analysis (EMSA), the purified proteins were electrophoresed in parallel with BSA standards. The gels were then stained with Coomassie Brilliant Blue R, scanned with a Chemi-Doc MP Imaging system (Bio-Rad) and the amount of recombinant protein determined by plotting a standard curve for the BSA (128). EMSA was carried out as described previously (55). Binding was carried out in 10% glycerol, 50 mM KCl, 5 mM MgCl2 and 1 mM DTT using fluorescent tagged DNA, nucleotides −30 to + 10 of the rat rDNA promoter, and no competitor. Gel electrophoresis was carried using Tris (25mM)/glycine (0.19M) buffer at pH 8.3. The gels were pre-electrophoresed for 30 min at 60V before loading and running in the dark. Western blots and shifts were visualized with a ChemDoc MP (Bio-Rad).

### Statistical Analysis

All experiments were reproduced at least three times with three technical replicates each time. Quantitative results that required comparisons between groups were subject to statistical analysis using two-tailed Student’s t test for two groups or one way ANOVA followed by Dunnett’s multiple comparison test to determine significant differences among more than two groups. Data met assumptions of the tests (*i.e*., normal distribution, similar variance). Normality of our data was determined via a D’Agostino-Pearson omnibus normality test and a Shapiro-Wilk normality test. To determine whether the variance differed between groups, an F test was performed when comparing two groups and a Bartlett’s and Brown-Forsythe test was used to compare the variance of more than two groups.

## Acknowledgements

This work was supported by GM069841 and HL077814 from NIH and HR15-166^1^ from OCAST awarded to L.I. Rothblum and funds from the University of Oklahoma. L.I.Rothblum is a member of the Stephenson Cancer Center.

## Footnotes

1 The content is solely the responsibility of the authors and does not necessarily represent the official views of the National Institutes of Health.

